# Single-cell analysis reveals diversity of tumor-associated macrophages and their interactions with T lymphocytes in glioblastoma

**DOI:** 10.1101/2023.08.15.553323

**Authors:** Sai Batchu, Khalid A. Hanafy, Navid Redjal, Saniya Godil, Ajith J Thomas

## Abstract

Glioblastoma (GBM) is an aggressive primary CNS malignancy and clinical outcomes have remained stagnant despite introduction of new treatments. Understanding the tumor microenvironment (TME) in which tumor associated macrophages (TAMs) interact with T cells has been of great interest. Although previous studies examining TAMs in GBM have shown that certain TAMs are associated with specific clinical and/or pathologic features, these studies used an outdated M1/M2 paradigm of macrophage polarization and failed to include the continuum of TAM states in GBM. Perhaps most significantly, the interactions of TAMs with T cells have yet to be fully explored. Our study uses single-cell RNA sequencing data from adult IDH-wildtype glioblastoma, with the primary aim of deciphering the cellular interactions of the 7 TAM subtypes with T cells in the GBM TME. Furthermore, the interactions discovered herein are compared to IDH-mutant astrocytoma, allowing for focus on the cellular ecosystem unique to GBM. The resulting ligand-receptor interactions, signaling sources, and global communication patterns discovered provide a framework for future studies to explore methods of leveraging the immune system for treating GBM.

## Introduction

Glioblastoma (GBM) is one of the most aggressive primary CNS malignancies. Despite advances in treatment protocols, clinical outcomes have not followed as evidenced by a 5-year survival rate near 5% [2]. Key obstacles remain, including an understanding of the tumor microenvironment (TME) which is heterogenous. Recent focus has shifted towards the cellular relations in this system, mainly the surrounding stromal cells of which macrophages are the most prominent and significant in disease progression [1, 3, 4].

Macrophages and microglia (local CNS macrophages derived from yolk sac) are both important components of the immune response. While microglia have self-renewal capacity and maintain brain homeostasis with minimal immune stimulation[5, 6], bone-marrow-derived macrophages infiltrate the brain only in pathological conditions [7]. Indeed, in GBM, the monocyte-derived macrophages may compose as much as 50% of the total mass [8]. However, unlike normal macrophages, these tumor-associated macrophages (TAMs) behave differently, altering the bridge between innate and adaptive immune responses and causing states of pathologic TAM-mediated immunosuppression.

A major component of the TAM mediated immunosuppression involves the lymphoid lineage, namely T cells. It has been observed TAMs may impede T cell functionality, promote T cell exhaustion, and impede T cell migration [9]. As macrophages exhibit high degrees of plasticity, they have historically been viewed in the M1 (pro-inflammatory) or M2 (anti-inflammatory) phenotypes, with the M1 macrophages associated with anti-tumorigenic effects. It is now accepted that this dichotomous nomenclature oversimplifies TAM plasticity, especially in a dynamic setting such as the TME of GBM [10–12]. To truly appreciate the cellular interactions of TAMs and T cells in GBM, a framework that includes a continuum of TAM activation states must be used.

Recently, 7 TAM subtypes were found to be preserved in almost all cancer types and were characterized using single-cell multi-omics technologies. These 7 TAM subsets, based on signature genes and enriched pathways, have been categorized as follows: interferon-primed TAMs (IFN-TAMs), immune regulatory TAMs (reg-TAMs), inflammatory cytokine-enriched TAMs (inflam-TAMs), lipid-associated TAMs (LA-TAMs), pro-angiogenic TAMs (angio-TAMs), resident-tissue macrophage-like TAMs (RTM-TAMs), and proliferating TAMs (prolif-TAMs) [12].

Although previous studies examining TAMs in GBM have shown that certain types are associated with certain clinical and/or pathologic features [1], these studies have used the outdated M1/M2 paradigm of macrophage polarization and failed to include the continuum of TAM states in GBM, and especially how they interact with T cells. Therefore, the present paper used single-cell RNA sequencing data from adult IDH-wildtype glioblastoma, with the primary aim of deciphering the cellular interactions of the 7 TAM subtypes with T cells inside the GBM TME. Furthermore, the interactions discovered herein are compared to IDH-mutated astrocytoma, allowing for focus on the cellular ecosystem unique to GBM. The resulting ligand-receptor interactions, signaling sources, and global communication patterns discovered provide a framework for future studies to explore methods in leveraging the immune system for treating GBM.

## Methods

### Data acquisition

Single cell RNA sequencing data matrices and metadata comprising myeloid and T cells from 7 newly diagnosed IDH wildtype GBM samples and 1 IDH mutant astrocytoma sample derived from a previous study was used for the present analysis [13]. Another dataset including immune cells from GBM samples was examined separately for in silico validation [14]. Additionally, previously published GBM spatial transcriptomics data (sample 269_T) was computationally examined to explore the spatial distribution of gene expression of certain cell type patterns in the tissue extracted from these samples with respect to the barcode-spot’s spatial dimensions [15]. ‘AUCell’ R package was used to annotate the TAM subpopulations from the original datasets. Gene signatures derived from previous literature [12] (Table 1) was used for TAM subpopulation classification. The R packages ‘celldex’ and ‘SingleR’ were used to classify the other immune cell populations, including T cells, into further subtypes using the ‘MonacoImmuneData’ reference index, which contains normalized expression values of bulk RNA-seq samples of 29 sorted immune cell populations, allowing for high resolution classification of immune cell transcriptomes [16].

**Table 1:**
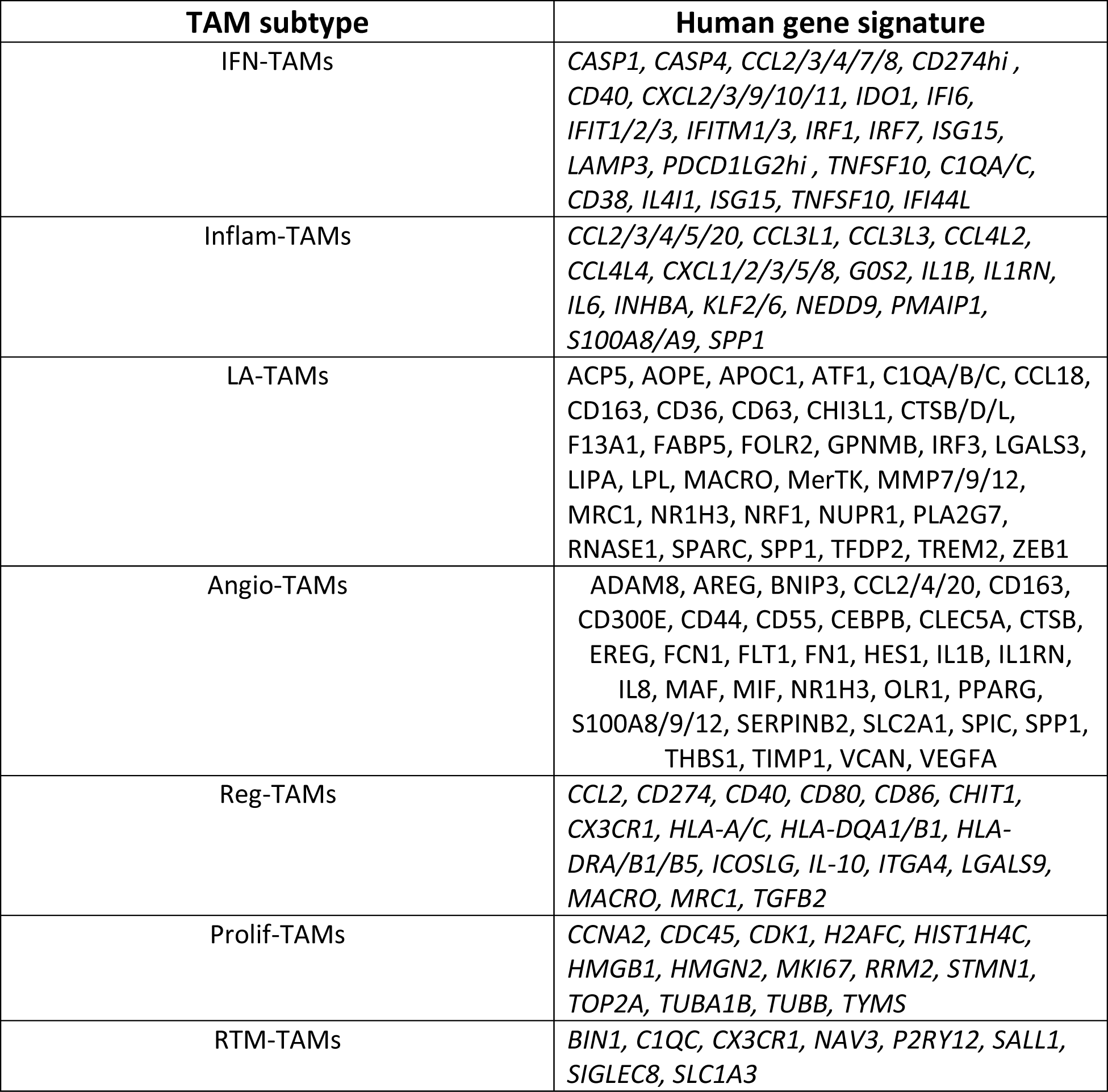
Gene signatures used to annotate TAMs in the current study. Adapted from Ma et al., 2022 [12].

### Ligand-receptor analysis

Cell communication analysis was based on CellChat [17]. The CellChatDB was used as the reference database of literature-supported ligand-receptor interactions, containing more than 1,900 validated and manually curated interactions including paracrine/autocrine secreted signaling interactions. Using this database of known ligand-receptor interactions, CellChat initially identifies differentially over-expressed receptors and ligands in each provided cell type. Each interaction is subsequently associated with a probability value, modelled by the law of mass action based on the average expression values of a receptor of one receiver cell type and the average expression values of the ligand of the counterpart cell type.

CellChat was run using ‘trimean’, a robust statistical method for calculating the average gene expression per cell type. This method assembles fewer interactions but performs better for predicting stronger interactions which is advantageous when selecting those interactions for further experimental validations. Lastly, the significance of these interactions is identified by randomly permuting the cell type labels and subsequently recalculating the interaction probability [17]. Comparisons between IDH wildtype and mutant for identifying up-regulated and down-regulated signaling was performed by comparing the communication probability between two datasets for each ligand-receptor pair and each pair of cell groups. Interactions with *p*-value < 0.05 were considered significant.

### Identifying major signaling sources and global communication patterns

To examine dominant senders and receivers within the interactome, the present study leveraged metrics derived from graph theory which were previously utilized for social network analysis [17, 18]. Specifically, weighted-directed networks and associated metrics, including out-degree and in-degree, were used to discover dominant intercellular interaction senders and receivers. In a weighted-directed network, the weights (i.e., strengths of interactions) are defined as the calculated communication probabilities based on the gene expression of ligands and receptors. Out-degree is calculated as the sum of these communication probabilities of outgoing signaling from an individual cell group. In-degree is computed as the sum of communication probabilities of incoming signaling to a cell group. These scores were used to identify the dominant cell senders and receivers, respectively, of intercellular signaling networks in the weighted-directed network [17]. Additionally, information flow was calculated for a signaling pathway by summing the communication probabilities amongst all pairs of cell groups in the predicted network [17, 19].

To identify communication patterns among all signaling pathways, non-negative matrix factorization was carried out, with the number of patterns based on the Cophenetic and Silhouette metrics, which measure the stability for a particular number of patterns via hierarchical clustering of the derived consensus matrix [17, 20]. Plainly, outgoing communication patterns show how sender cell types coordinate with each other and how they work with certain signaling pathways to support communication. Correspondingly, incoming patterns reveal how target cell types (i.e., cells receiving signals) coordinate with each other to respond to incoming signaling.

## Results

Using dimensionality reduction, the TAMs from GBM and IDH mutant glioma showed distinct clustering profiles as well as areas of overlap (Fig. 1A). All seven types of TAMs were found in both types of gliomas (Fig. 1B). IDH wildtype GBM harbored more LA TAMs, inflammatory TAMs, proliferative TAMs, but had less RTM microglia-like TAMs.

**Figure 1:**
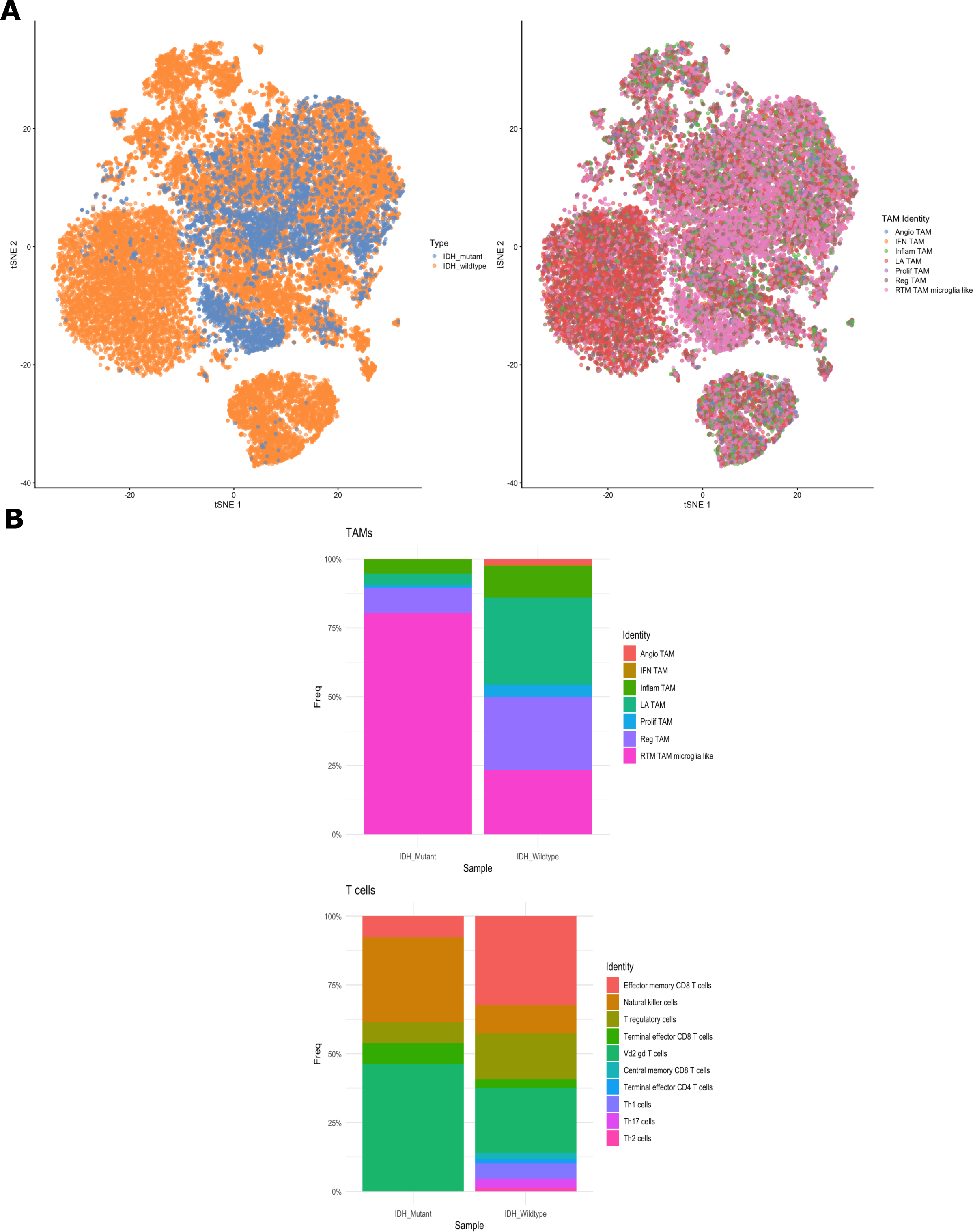
**(A)** t-SNE plot of TAMs **(B)** Stacked bar plot displaying frequency of cell types in both IDH wildtype and mutant tumors.

To predict significant ligand-receptor interactions between TAMs and T cells, a comprehensive dataset including single cell gene expression data from both IDH wildtype primary GBM and IDH mutant gliomas was used [13]. To understand which cell types are dominant senders (i.e., cell types sending signals) and which are dominant receivers (i.e., cell types receiving signals) with respect to each ligand-receptor interaction analyzed, incoming and outgoing interactions strengths (weights) were calculated for both GBM and glioma (Fig. 2A). In both IDH wildtype and mutant, angio-TAMs displayed robust incoming and outgoing signaling strength. In the IDH wildtype, all seven TAM types displayed strong outgoing interaction strength compared to IDH mutant, where only Inflam TAMs and LA TAMs showed strong outgoing interaction strength.

**Figure 2:**
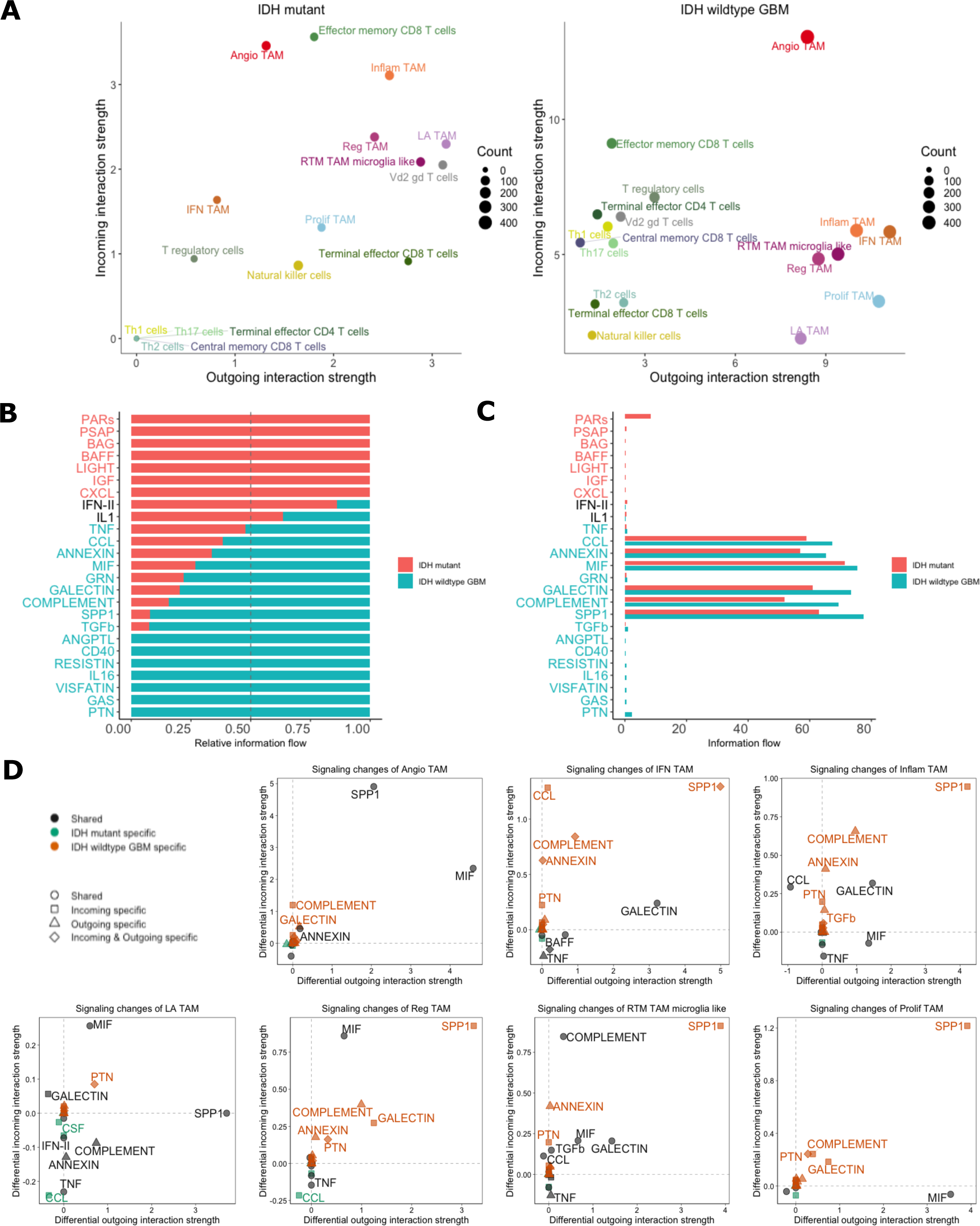
**(A)** Scatterplot displaying dominant senders and receiver cell type. The total outgoing and incoming communication probability associated with each cell group is displayed on x-axis or y-axis, respectively. Count is displayed by dot size which is proportional to the number of predicted signaling links associated with each cell group. **(B)** Stacked bar plot and unstacked bar plot **(C)** showing conserved and context specific signaling pathways by comparing information flow for each signaling pathway. **(D)** Scatterplot showing specific signaling changes of TAM populations. Positive values indicate increase in IDH wildtype GBM compared to IDH mutant astrocytoma.

Next, we then sought to identify those signaling pathways that were conserved and those that were specific to GBM by calculating the information flow within signaling pathways, defined as the sum of communication probabilities amongst the total pairs of cell groups in the network (i.e., the amount of activity occurring in the pathway) (Fig. 2B). Compared to the IDH mutant astrocytoma, GBM exhibited increased information flow in multiple pathways including GALECTIN, COMPLEMENT, MIF, SPP, and PTN. When considering the absolute information flow in each pathway (Fig. 2C), greater information flow was observed in CCL, ANNEXIN, MIF, GALECTIN, COMPLEMENT, and SPP1 pathways. These flow states imply that although IDH wildtype GBM has increased activity in these pathways relative to IDH mutant astrocytoma, both types of gliomas have substantial activity occurring within these signaling pathways.

Since IDH-wildtype GBM displayed prominent activity of TAMs (Fig. 2A) and displayed specificity for certain pathways (Fig. 2B, C), we then sought to identify the specific signaling changes of the different TAM populations between GBM and IDH-mutant (Fig. 2D). Angio-TAMs and LA TAMs were observed to have shared signaling amongst both glioma types in SPP1 and MIF pathways. However, the other five TAM subtypes showed increased signaling in SPP1 specific to GBM (Fig. 2D). Overall, the seven TAM populations were observed to be contributing in an increased capacity to pathways such as SPP1, COMPLEMENT, GALECTIN and PTN compared to their counterparts in IDH mutant astrocytoma.

Further examination of each pathway yielded numerous significant ligand-receptor interactions (Fig. 3A, B). All types of TAMs exhibited MIF and SPP1 binding interactions with ligands on T cells such as CD44 and ITGA4+ITGB1 (Fig. 3A). Conversely, T cell ligands interacting with receptors on TAMs commonly included MIF-(CD74+CD44) and ANXA1-FPR1 (Fig 3A). To further verify these interactions, another dataset that included was analyzed independently in a similar fashion [14]. The ligand-receptor interactions were corroborated by the second dataset, as interactions including SPP1, MIF, LGALS9 between TAMs to T cells were found to be significant (Fig 4A, 4B). When deciphering the ligand-receptor interactions in the context of cell-to-cell contact (Fig. 5), numerous significant interactions between HLA class I molecules and CD8 were observed in GBM. Such HLA class I molecules were indicated to be on TAMs, including HLA-A, B, C, E, and F. Additional interactions from TAMs to effector CD8 T cells such as CD99-CD99 were also seen. Activated leukocyte cell adhesion molecule (ALCAM) interacting with CD6 on T cells were witnessed to be significant in most of the interactions from TAMs to T cells (Fig. 5).

**Figure 3:**
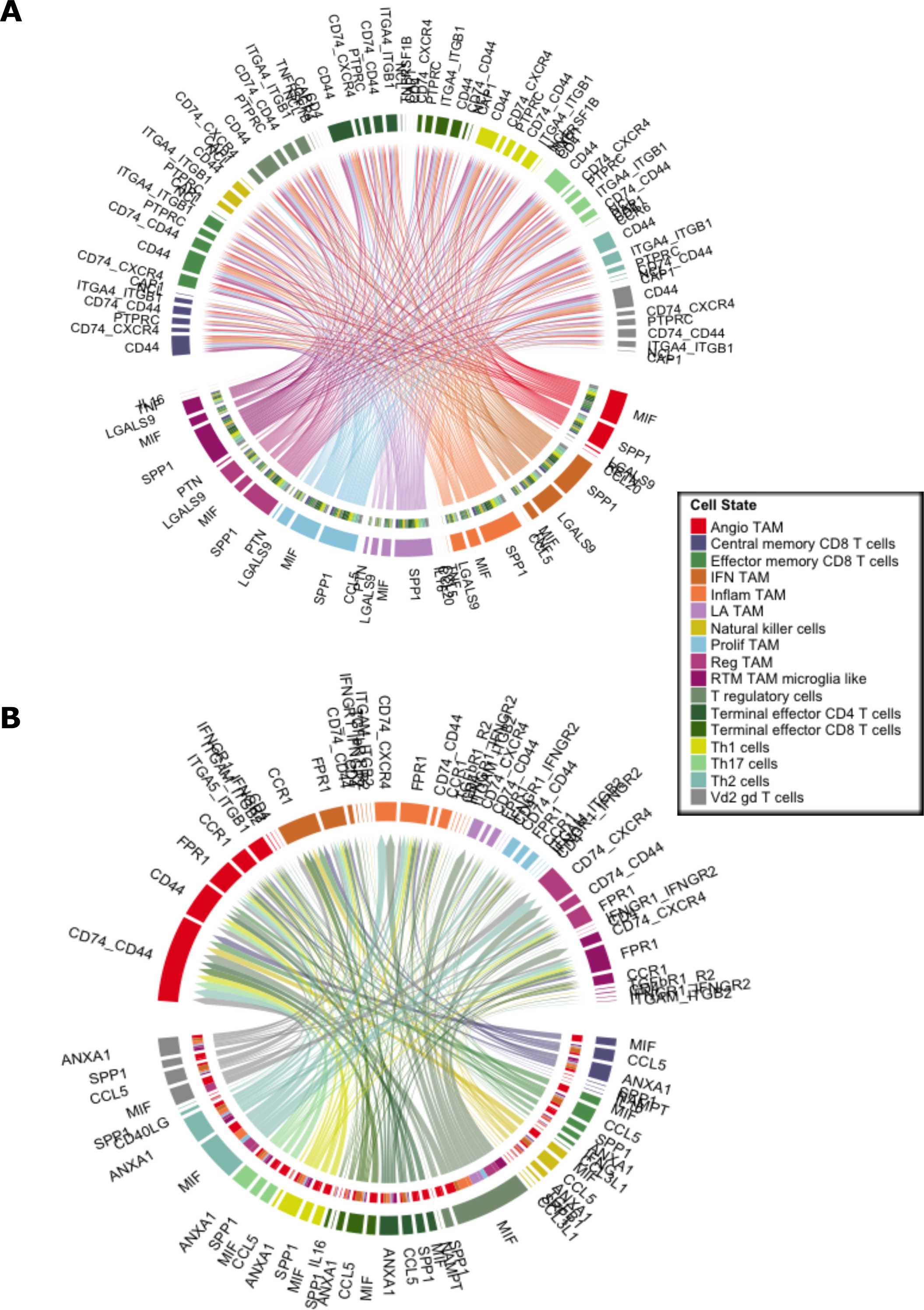
**(A)** Chord diagram visualizing cell-cell communication for significant ligand-receptor interactions from TAM populations (senders, bottom arc) to T cell populations (receivers, top arc). **(B)** Chord diagram visualizing significant ligand-receptor interactions from T cell populations (senders, bottom arc) to TAM populations (receivers, top arc).

**Figure 4:**
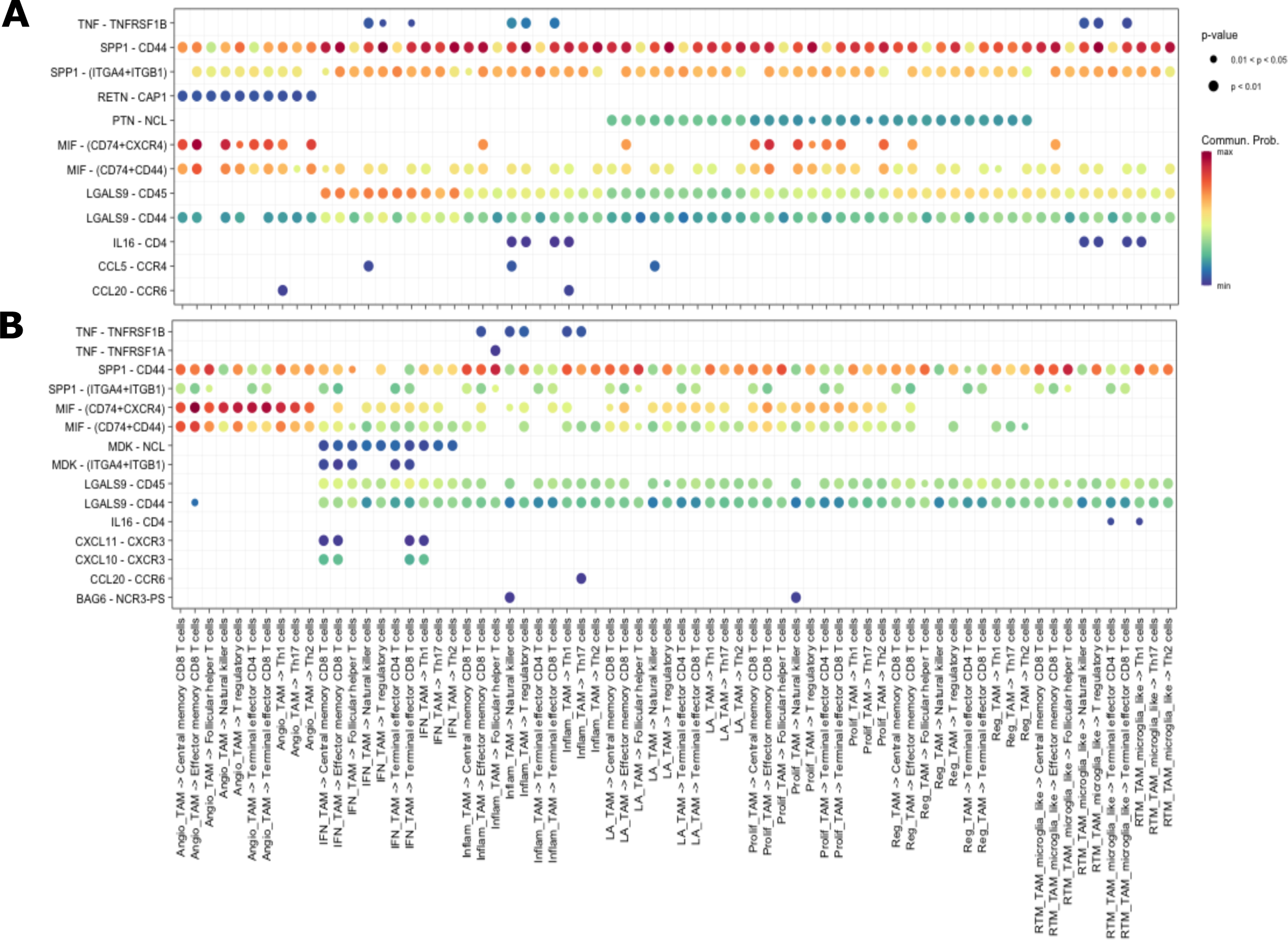
Bubble plot showing all the significant secreted signaling interactions (L-R pairs) (y-axis) from TAM populations (senders) to T cell populations (receivers) (x-axis). The dot color and size represent the calculated communication probability (represents interaction strength) and p-values (represents statistical significance). Non-significant interactions represented by blank spaces. **(A)** Abdelfattah et al., 2022 dataset. **(B)** Additional independent analysis using Pombo Antunes et al., 2021 dataset.

**Figure 5:**
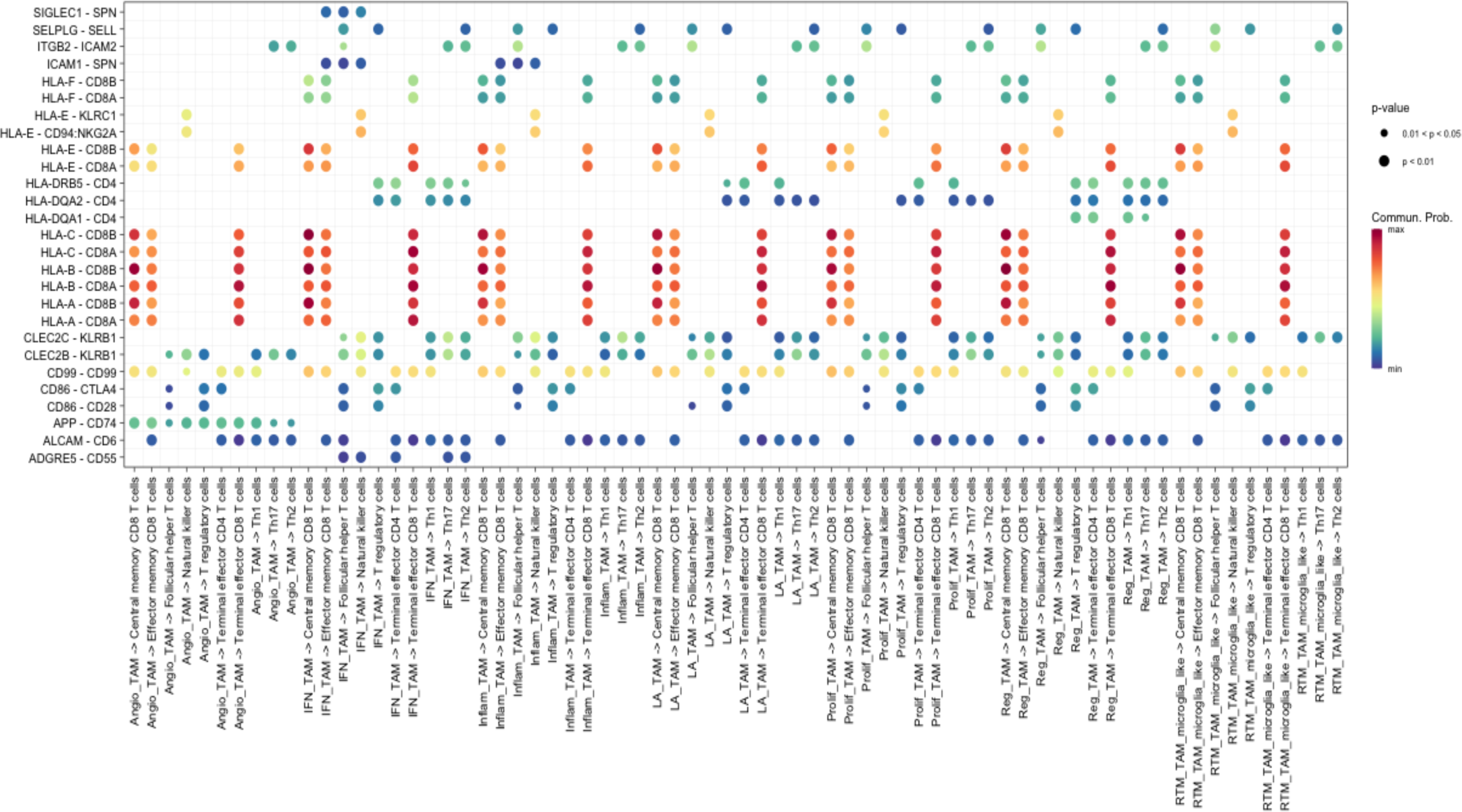
Same as Figure 4 but for cell-to-cell contact interactions.

Along with understanding ligand-receptor interactions in the tumor microenvironment, it is crucial to analyze how different cell types coordinate with each other in terms of signaling. To address this, a pattern recognition methodology was employed to identify global communication patterns of TAMs and T cells (see Methods). These communication patterns can be thought of as assigning cell types to signaling pathways for either outgoing signaling (considering cells as senders) or incoming signaling (considering cells as receivers) (Fig. 6A, B). The outgoing cell signaling patterns were broadly identified into 3 distinct categories. Most TAMs contribute to pattern 1, while pattern 2 was mainly contributed by angio-TAMs and T regulatory cells (Fig. 6A). Pattern 1 was mainly composed by pathways involving TNF, GALECTIN, SPP1, and COMPLEMENT while pattern 2 mainly composed of RESISTIN and ANGPTL signaling (Fig. 6A). For the incoming signaling pathways in IDH wildtype GBM, 3 discernable patterns also emerged (Fig. 6B). Interestingly, angio-TAMs displayed pattern 3, as opposed to the other TAMs which displayed a pattern 1 incoming signaling. Pattern 3 is mostly contributed by VISFATIN, IL-1, ANGPTL and CD40.

**Figure 6:**
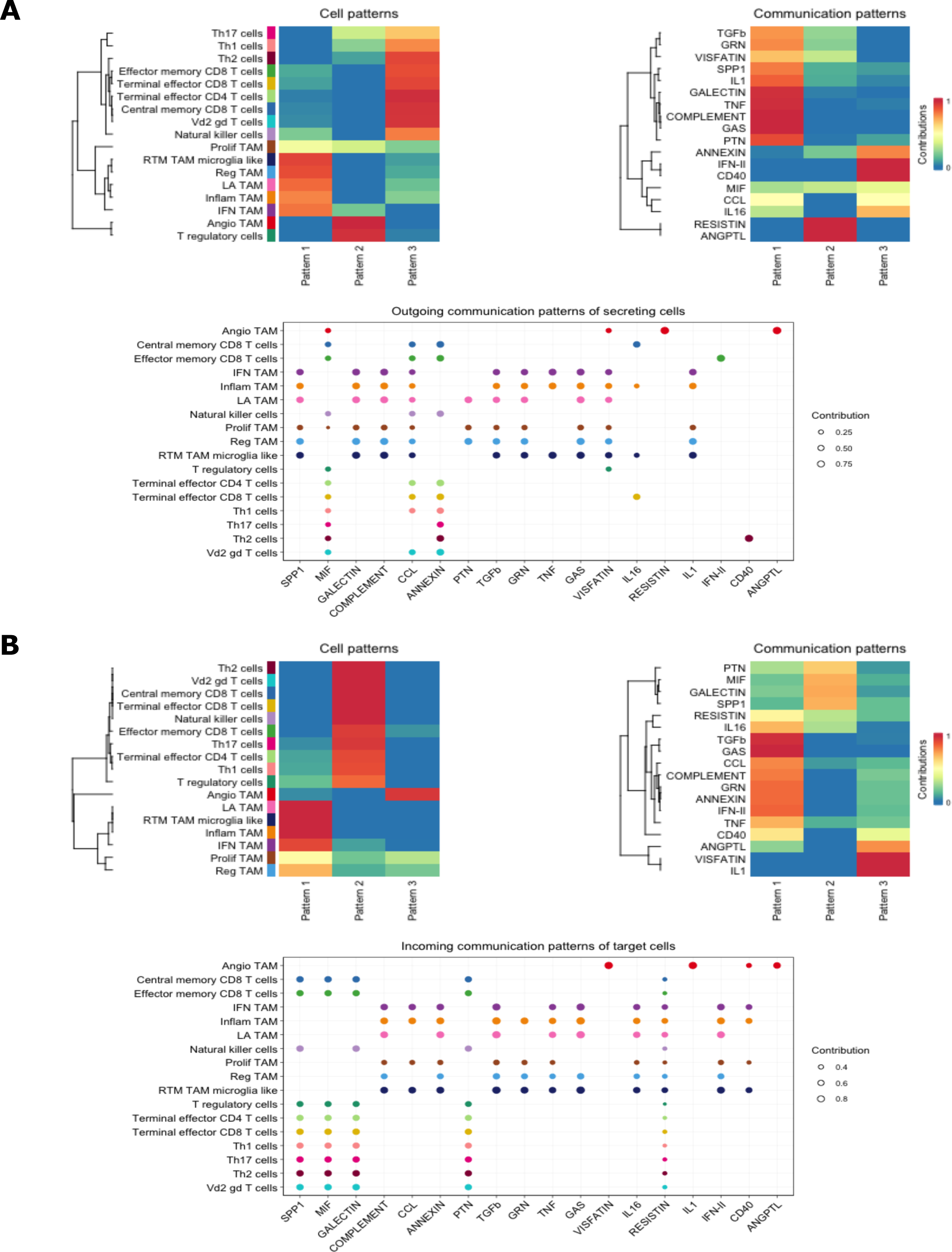
Global communications analysis for **(A)** outgoing and **(B)** incoming communication patterns. Cell patterns heatmap display which cell types contribute to which type of pattern. Communication patterns heatmap displays which signaling pathways contribute to the pattern. The dot plots show how each cell group contributed to the signaling pathways, where the dot size is proportional to the contribution score to show association between cell group and their enriched signaling pathways. A higher contribution score implies the signaling pathway is more enriched in the respective cell type.

These communication results imply that TAMs in GBM may depend on numerous signaling networks that may overlap in function while certain TAMs, such as angio-TAMs, rely on fewer and more heterogenous communication patterns. Additionally, leveraging these incoming and outgoing patterns may provide insight into autocrine vs. paracrine pathways for a given cell type in the GBM cellular landscape.

Given this interplay amongst TAMs and T cells, especially within the context of T cell exhaustion in GBM [21], a better understanding of the spatial interactions of these cell types should be explored. First, examining genes related to T cell exhaustion revealed an increased expression in T cells within GBM (Fig. 7). Specifically, T regulatory lymphocytes in GBM exhibited increased expression of *ICOS, CTLA4, TIGIT, IL2RA,* and *IL10RA* (Fig. 7). To evaluate further, spatial transcriptomic data from a GBM dataset [15] was examined by transposing the gene signatures of T cell exhaustion and the TAMs analyzed in the present study (Fig. 8). The T cell signature used included persistent expression of multiple inhibitory receptors, such as *PD-1, LAG-3, TIM-3, and CTLA-4,* and TIGIT [21]. The spatial plots displayed a prominent overlap amongst TAM signatures and the T cell exhaustion gene signature, with a higher density and expression of T cell exhaustion related genes in spaces occupied by TAM signatures (Fig. 8)

**Figure 7:**
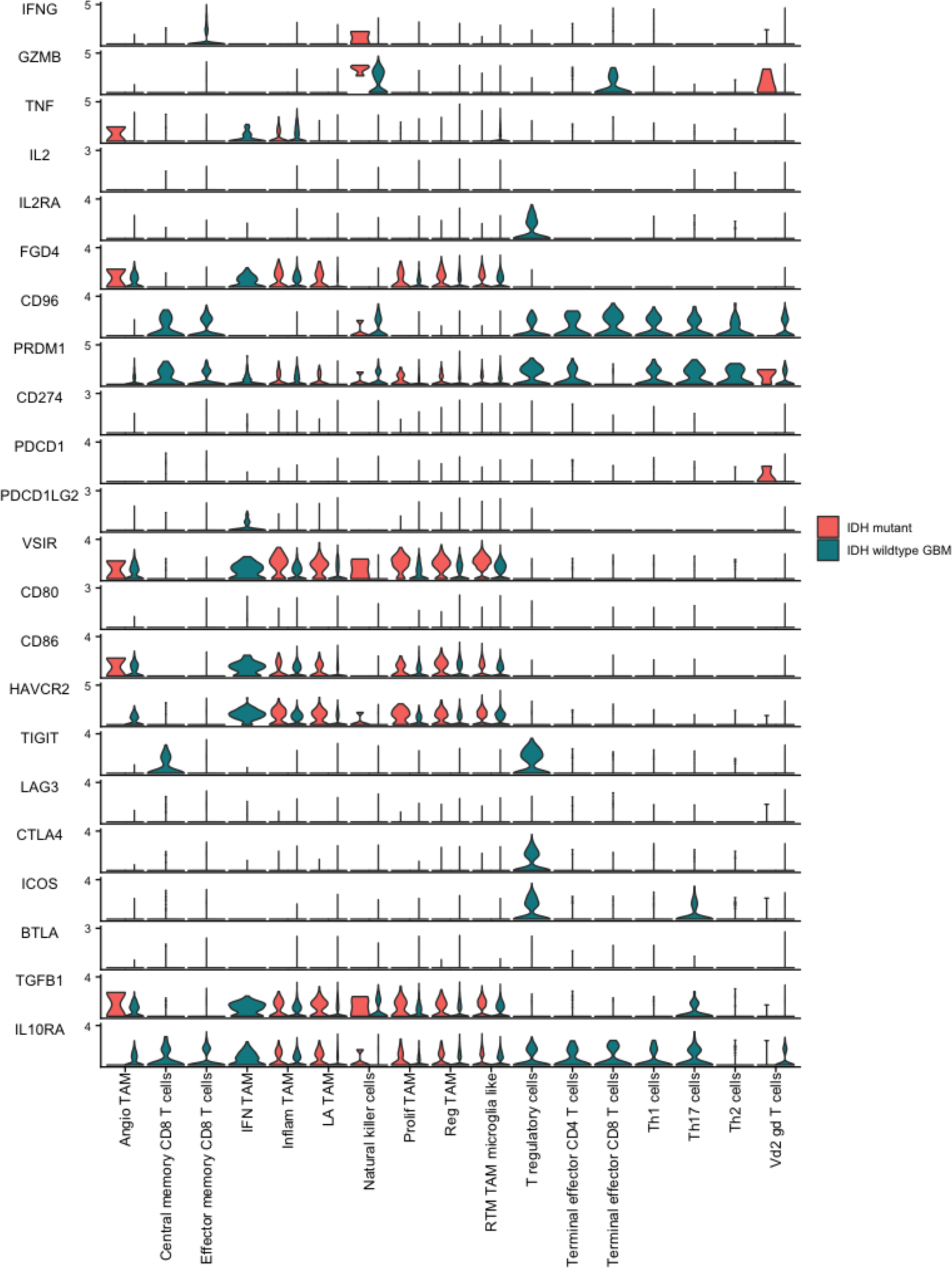
Violin plot of gene expression values in TAMs and T cells between GBM and IDH mutant glioma.

**Figure 8:**
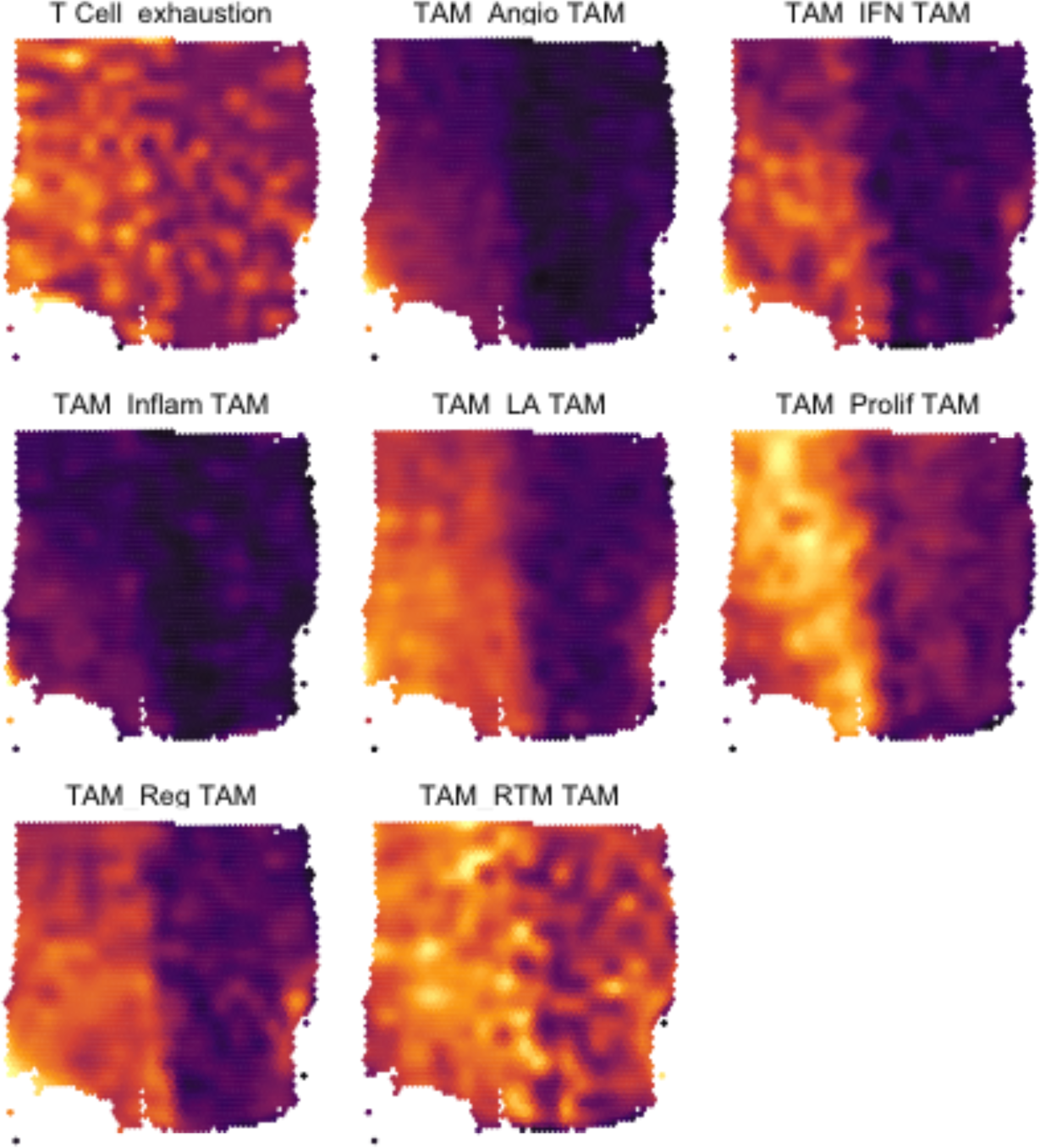
Surface plot displaying the expression signatures of the T cell exhaustion cell state and seven TAM types derived from spatial transcriptomics. Brighter areas correspond to higher average expression of genes comprising the signature. Color intensities are exclusive to each plot.

## Discussion

### IDH mutant gliomas and GBM have distinct immune profiles diverse repertoire of TAMs

All seven subsets of TAMs were found in both IDH wildtype GBM and mutant gliomas. Angio-TAMs have an angiogenic signature associated with significant immune suppression and tumor cell proliferation. Though found in proportionally smaller numbers, they were observed to be the dominant receivers and senders in the population communicating with the lymphocyte population. Angio-TAMs were rarely encountered in IDH mutant gliomas. Both IDH mutant and IDH WT gliomas have increased representation of myeloid cells with lymphocyte population increased in the IDH WT tumors [4]. RTM-TAMs formed the largest component of TAMs in the IDH mutant variety. RTM-TAMs are very similar to normal RTMs and a putative role for them in tumor invasiveness in lung cancer has been suggested [22]. LA-TAMs and proliferative TAMs were also seen in larger numbers in the WT high grade gliomas. LA-TAMs express more lipid related and oxidative phosphorylation genes which include *APOC1*, *APOE*, *ACP5* and *FABP5*.Lipid catabolism in macrophages is associated with IL-4 associated pathways and anti-inflammatory phenotype[23]. Inflammatory TAMs have an inflammatory signature and express inflammatory cytokines including IL1B, CXCL1/2/3/8, CCL3, and CCL3L1. Inflammatory TAMs are known to play an active role in the recruitment of immune cells including granulocytes into the TME. Regulatory TAMs, which were found more often in GBM, seemed to have a profile consistent with microglia and appeared to play a more quiescent role in activating lymphocytes. This was corroborated by Venteicher et al. who demonstrated microglia in IDH mutant glioma TME were associated with a less malignant phenotype [24].

Effector memory CD8 T cells were more often encountered in WT glioma while NK cells were more common in IDH mutant glioma. Th1, Th2 and Th 17 CD4+ T cells were hardly seen in IDH mutant gliomas, compared to IDH-WT tumors. Vd2 gd T cells were present abundantly in both groups.

### TAMs are associated with T cell and NK cell exhaustion in the TME

T lymphocyte exhaustion is a well-known phenomenon in GBM with cells expressing several inhibitory checkpoint receptors including PD-1, LAG-3 and CTLA-4 [25]. Leveraging spatial transcriptomics revealed that signatures of all seven TAM subtypes were associated with a strong expression of genes related to T cell exhaustion. Ravi et al. previously identified a subset of TAMs that secreted IL-10 and HMOX1+ and contributing to T cell exhaustion in glioblastoma [26]. These cells were characterized by the expression of *CD163, CCL4, APOE* and *HLA –DRE*. In our dataset, they would map to angio-TAMs, LA TAMS and Inflam TAMs. The TAMs were observed to produce an immunosuppressive environment by secreting ligands such as MIF, Galectin 9 and SPP1 and affecting cell-to-cell contact and are described in subsequent sections.

*CD86, HAVCR2 (TIM3), TGFB1* were significantly expressed in TAMs. *TIGIT* was upregulated in central memory CD8 T cells. *ICOS (CD278), CTLA4* and *TIGIT* were robustly expressed in T regs*. IL-10 RA*, which codes for the IL-10 receptor, is well-known to induce immunosuppression. It was strongly expressed in all the immune cells consistent with prior studies showing that IL-10 contributes to the immunosuppressed environment in gliomas [26].

Transforming growth factor-beta (TGF-β), which promotes an immunosuppressive milieu, was also seen to be expressed by all TAMs in secreted signaling interactions (Fig. 7). *PRDM1*, known to regulate PD-L1 levels resulting in CD8+ T cell exhaustion, was also expressed in all immune cells examined [27].

*CD96*, an immune checkpoint ligand, was strongly expressed in T and NK lymphocytes within GBM samples. VSIR, an important checkpoint modulator in AML, was highly expressed in both IDH mutant and GBM TAM populations [28, 29]. *FGD4*, albeit not well studied, was also well represented in the same group of cells [30, 31] (Fig. 7). Furthermore, TAMs interact with various immune cells within the tumor microenvironment through other mechanisms which include expression of Indoleamine 2,3-dioxygenase (IDO) and production of Arg1 which is beyond the scope of the current analysis.

### TAMs in GBM secrete a common group of ligands which are immunomodulatory

Immune cells in the TME of GBM secrete a variety of immunomodulatory molecules [21, 26]. During the present analysis, certain prominent signaling common amongst TAMs and T cells in GBM were revealed, including galectin, complement, MIF, SPP and PTN signaling pathways (Fig. 2-4). In addition, when considering the absolute information flow in each pathway, CCL and ANNEXIN also assumed importance (Fig. 2). All these signaling pathways account for significant immune suppression and support tumor proliferation.

Regardless of the TAM subtype, SPP1 (Osteopontin) seemed to be very significant in outgoing and incoming signaling (Fig. 2B-D). SPP1 suppresses T cell function through the IRF8-OPN-CD 44 axis [32]. MIF (Macrophage Inhibitory Factor) also can act to reduce CD8+ Cytotoxic lymphocyte (CTL) activity [33, 34]. We found that there was significant upregulation of the MIF ligand - CD74/CD44 pair, which can recruit macrophages and foster an immunosuppressive niche. Blocking this interaction in TAMs and dendritic cells in melanoma showed decrease in immunosuppression [35]. MIF also interacts with immune checkpoint molecules like PD-L1, contributing to T cell exhaustion and immune evasion while suppressing cytotoxic T cells and NK cells[33]. Complement secreted by TAMs is increasingly seen as immunosuppressive and promotes tumor growth. This immunosuppressive effect is mediated through multiple pathways which include the increased expression of molecules such as PDL-1, IL-10, Arg-1, and TGF-β1[36].

Our results also revealed significant expression of interactions involving galectins and the galectin signaling pathways (Fig. 2-4, 6). Galectin 1 is highly expressed in glioblastoma stem cells and is important for immunomodulation and angiogenesis[31–33]. Galectin 9 is associated with T cell exhaustion in the tumor environment[38, 39]. The binding of TIM3 to its ligand, galectin-9 induces T cell apoptosis [33]. Galectin binds to the CD45 receptor as shown in the ligand interaction and CTLA 4 to reduce T cell proliferation and induce T cell apoptosis [43]. Galectin is also involved in the formation and stability of regulatory T cells (suppressors of antitumor immunity) through the CD44 receptor in association with TGF-β. The CD44 receptor is crucial in mediating the action of SPP1, Galectin and MIF. CD44 is glycoprotein which is used as a marker for cancer stem cells (CSC) and has significant role in the maintenance and progression of solid tumors [40]. CD44 is also involved in the positive regulation of the immune checkpoint molecule PD-L1, macrophage infiltration of tumor and contributes to the antitumor activity of TAMs[41].

The annexin pathway is also revealed to be crucial in the secreted signaling interactions between TAMs and T cells (Fig. 2-4, 6). Annexin 1 is expressed in glioblastoma and known to promote tumor growth while fostering an immunosuppressive environment [42, 43]. Treatment of macrophages with AnxA1 results in decreased iNOS expression, increased IL-10 levels and decreased IL-12 mRNA levels and an anti-inflammatory phenotype [42]. CCL (chemokine ligands) secreted can also produce an environment that is favorable to tumor progression or result in anti-tumor activity. The recruitment of monocytes from peripheral blood and differentiation into TAMs are dependent on the CCR5-CCL5 and CCR2-CCL2 chemokine axes. TAMs also secrete CCL17, CCL22, and CCL18 which attract regulatory T cells [44]. Pleiotrophin (PTN), another significant result in our analyses, is known to be secreted by TAMs and enhance glioblastoma cell proliferation through its receptor PTPRZ1 expressed on glioma stem cells (GSCs)[45]. PTN also contributes to angiogenesis in gliomas and increased expression of PTN is associated with poor survival in glioma patients [46].

Visfatin/NAMPT, originally isolated from peripheral lymphocytes, can induce monocytes to transform into TAMs with CD163 expression. Indeed, angio-TAMs express CD163 strongly on their surface (Table 1) [12]. These TAMs also secrete the chemokine CXCL1 which are known to increase invasion and migration of cancer cells along with angiopoietins that increase angiogenesis and promotes tumor growth in glioblastomas [47, 48].

Angiopoietin-like proteins (ANGPTLs) are structurally like angiopoietins. We found multiple interactions involving these molecules in the current analysis. Angiopoietin-like 2 (ANGPTL2) fosters the conversion of TAMs into a less inflammatory subset in NSLC. It also correlates positively with TAM infiltration and a poor prognosis[47, 49]. Another family member, angiopoietin-like 4 (ANGPTL4) blunts the polarization of macrophages toward the proinflammatory phenotype and decreases immune surveillance in tumor progression by downregulating CD8 T cell activation [50]. Resistin was another key mediator in our analyses. Mostly expressed by macrophages in humans, it has potent pro-inflammatory properties and is shown to promote cancer progression while inducing expansion of regulatory T cells which have an immunosuppressive role [51, 52].

### TAMs in WT modulate T and NK lymphocytes through cell-to-cell contact

TAMs also communicate with lymphocytes through cell-to-cell contact (Fig. 5). Since the TAMs provide an immunosuppressive environment, it could be assumed that these interactions are inhibitory. Surprisingly the most prominent of these interactions center around the MHC Class I molecules which reiterates recent insights into how TAMs regulate the immune environment [53, 54]. Our results displayed that HLA – A, B, C and E interact with CD8 receptors significantly on central memory, effector memory, and terminal effector T cells. Indeed, HLA – A, B,C have been shown to negatively impact the function of NK cells and cytotoxic T lymphocytes through engagement with LILRs and lymphocytic KIR receptors resulting in immunosuppression, anergy, and T cell exhaustion [53, 54]. HLA-E on the surface of TAMs also bind to the inhibitory receptor CD94/NKG2A found on both NK cells and cytotoxic lymphocytes [55]. With regards to MHC class II molecules, HLA – DRB5 and CD4+ cells also interacted in cell-to-cell contact (Fig. 5). HLA-DRB5 has been found to be restrictive in presenting tumor antigens on CD 4 + cells [56].

This interaction is of critical importance because it forms the frontline against cancer propagation. Unfortunately, in advanced cancers, such as GBM, the TAMs seem to be tumorigenic with regards to their immunosuppressive role in the tumor microenvironment and T cell activation. While the role of Leukocyte Immunoglobulin-Like Receptors (LILRs) and Killer cell Immunoglobulin-like Receptors (KIRs) were initially evolved to quell an overactive immune response; tumors have hijacked the response for immune evasion and T cell tolerance, allowing the tumor cells to grow unabated.

Other interactions such as those involving CLEC2D and receptor KLRB1 were also seen. Cytotoxic T cells in glioblastoma express the NK lymphocytic gene *KLRB1*. TAMs express the ligand CLEC2D on the surface which serves as an inhibitor for KLRB1 receptor to suppress their tumor lytic function [57]. The homophilic CD99-CD99 interaction was demonstrated by most TAMs in our analyses. Previous studies show CD99 ligation presents an apoptotic feature in developing T lymphocytes, with increased CD99 expression associated with immune adaptation dominated by TAMs in gliomas [58]. The TAMs also interacted with lymphocytes through expression of CD86, which acts as the dominant ligand for CD4+FoxP3+ regulatory T cell survival and proliferation, interacting with CD28 and CTLA4 receptors on the regulatory T cells [59].

### Future directions

The immune landscape in glioblastoma is complex and most importantly not static. It evolves along with the progression of tumor. Immunosuppressive macrophages dominate towards the later stages of glioma evolution. We have identified several subsets of macrophages among which angiogenic TAMs play a very significant role. The immunosuppressive molecules of galectin, SPP1, MIF and complement impede the T cell response. In addition to the immunosuppressive role in the TME, the present study supports the view that TAMs directly facilitate glioma cell proliferation through its secreted ligands such as galectin and SPP1 along with direct cell-to-cell contact mechanisms.

Immunotherapeutic agents including checkpoint inhibitors and CAR T cell therapies have not shown encouraging utility in GBM, which is consistent with the immunologically ‘cold’ nature of GBM implying that the microenvironment is pervaded by immunosuppressive TAMs [60, 61]. TAMs seem to be critical in the trifecta of reducing macrophage activation, inhibiting cytolytic function in NK and cytotoxic cells, and reverting T cells to immunosuppression, Targeting TAMs in cancer, perhaps through “re-education into a pro-inflammatory state is critical to explore if future immunotherapies should hope to have any substantial effect in outcomes.

